# Plasma membrane transbilayer asymmetry of PI(4,5)P_2_ drives unconventional secretion of Fibroblast Growth Factor 2

**DOI:** 10.1101/2025.04.19.649664

**Authors:** Manpreet Kaur, Fabio Lolicato, Walter Nickel

## Abstract

Unconventional secretion of Fibroblast Growth Factor 2 (FGF2) is based upon direct self-translocation across the plasma membrane, a process that involves the transient formation of a lipidic membrane pore. The opening of this pore is triggered by PI(4,5)P_2_-dependent oligomerization of FGF2 at the inner plasma membrane leaflet. Subsequently, FGF2 oligomers populating the pore are captured by membrane-proximal heparan sulfate chains of Glypican-1 (GPC1), resulting in translocation of FGF2 to cell surfaces. PI(4,5)P_2_ is a highly negatively charged membrane lipid that is exclusively localized at the inner plasma membrane leaflet. Therefore, local accumulation of PI(4,5)P_2_ triggered by FGF2 oligomerization at the inner plasma membrane leaflet produces a steep and spatially restricted electrochemical gradient across the plasma membrane. Furthermore, PI(4,5)P_2_ has a wedge-like shape, turning it into a non-bilayer lipid that destabilizes membranes at high local concentrations. Here we demonstrate that an asymmetric distribution of PI(4,5)P_2_ across the leaflets of synthetic lipid bilayers accelerates the opening of FGF2-induced membrane pores. Consistently, we find unconventional secretion of FGF2 from cells to be inhibited under conditions compromising the native transbilayer asymmetry of PI(4,5)P_2_ of plasma membranes. We propose the asymmetric distribution of PI(4,5)P_2_ to lower the free energy required to transform the lipid bilayer into a lipidic membrane pore. Furthermore, in the proximity of FGF2 membrane translocation sites, PI(4,5)P_2_ in the outer plasma membrane leaflet could potentially repel negatively charged heparan sulfates chains compromising the function of GPC1 in FGF2 translocation into the extracellular space. Thus, transbilayer asymmetry of PI(4,5)P_2_ is a key parameter enabling fast kinetics of FGF2 membrane translocation into the extracellular space.

## Introduction

Fibroblast Growth Factor 2 (FGF2) belongs to a subclass of secretory proteins that do not travel along the ER/Golgi-dependent secretory route to reach the extracellular space. Such processes have collectively been termed ‘unconventional protein secretion’ (UPS) (Rabouille, 2017; Sparn et al., 2022b; Neel et al., 2024). Several alternative secretory routes have been discovered with FGF2 being the prime example of a UPS type I cargo protein that is secreted by direct translocation across the plasma membrane (Dimou and Nickel, 2018; Pallotta and Nickel, 2020). This process is initiated by sequential interactions of FGF2 with molecular components associated with the inner plasma membrane leaflet with (i) the Na,K-ATPase, (ii) Tec kinase and (iii) the phosphoinositide PI(4,5)P_2_ (Temmerman et al., 2008; Ebert et al., 2010; Steringer et al., 2012; Zacherl et al., 2015; La Venuta et al., 2016; Legrand et al., 2020). The Na,K-ATPase and Tec kinase have been proposed to represent auxiliary factors promoting efficient secretion by accumulating FGF2 at the inner plasma membrane leaflet. In addition, Tec kinase may play a role in enhancing FGF2 secretion in a genuine cancer context with high levels of PI(3,4,5)P_3_, resulting in enhanced membrane recruitment of Tec kinase. By contrast, PI(4,5)P_2_ has been shown to be an essential component of the core machinery of unconventional secretion of FGF2. The interaction with PI(4,5)P_2_ causes FGF2 to oligomerize on the surface of the inner plasma membrane leaflet, resulting in the opening of a lipidic membrane pore with a toroidal architecture through which FGF2 translocates into the extracellular space (Steringer et al., 2017; Lolicato et al., 2022; Lolicato et al., 2024). This process is driven by the cell surface heparan sulfate proteoglycan GPC1 that, by direct competition for the binding site of PI(4,5)P_2_ in FGF2, captures and disassembles FGF2 oligomers at the outer plasma membrane leaflet (Zehe et al., 2006; Sparn et al., 2022a), making FGF2 available for entering ternary complexes with heparan sulfate chains and high affinity FGF receptors for autocrine and paracrine signaling (Akl et al., 2016). A number of UPS type I cargo proteins have been identified (Malhotra, 2013; Rabouille, 2017; Dimou and Nickel, 2018; Pallotta and Nickel, 2020; Balmer and Faso, 2021; Sparn et al., 2022b), including HIV-Tat and Engrailed2 that are secreted in a PI(4,5)P_2_ and/or heparan sulfate dependent manner (Rayne et al., 2010; Debaisieux et al., 2012; Amblard et al., 2020; Joliot and Prochiantz, 2022).

The mechanism underlying unconventional secretion of FGF2 has been reconstituted with purified components using an inside-out topology system based on giant unilamellar vesicles (GUVs) (Steringer et al., 2017). The minimal machinery driving FGF2 membrane translocation could be identified with PI(4,5)P_2_ and heparin—as a surrogate of heparan sulfates—on opposing sides of the membrane. Membrane translocation could only be observed with FGF2 containing all *cis* elements required for unconventional secretion from cells such as the PI(4,5)P_2_ binding pocket and C95 required for oligomerization (Zehe et al., 2006; Temmerman et al., 2008; Müller et al., 2015; Lolicato et al., 2024). FGF2 membrane translocation into the lumen of GUVs could be observed at a time scale of minutes (Steringer et al., 2012; Steringer et al., 2017). In addition to *in vitro* approaches, individual events of FGF2 membrane translocation across plasma membranes have been visualized in intact cells, using real-time TIRF microscopy with single molecule resolution (Dimou et al., 2019). This study revealed extremely fast kinetics of FGF2 membrane translocation in a cellular context with an average time interval for individual events of about 200 milliseconds. Thus, a substantial difference in the kinetics of FGF2 membrane translocation could be observed between *in vitro* experiments under defined conditions and FGF2 secretion from intact cells (Sparn et al., 2022b).

Several parameters may play fundamental roles in governing the thermodynamic principles underlying fast FGF2 membrane translocation kinetics observed in living cells (Dimou et al., 2019). Among them are the Na,K-ATPase as a component to accumulate FGF2 at the inner plasma membrane leaflet of intact cells, presumably in close spatial proximity to GPC1 (Lolicato and Nickel, 2022). In addition, in cells, oxidative dimerization of FGF2 is likely to be mediated by an enzymatic mechanism, driving fast kinetics of PI(4,5)P_2_-dependent oligomerization. A third potential factor is the asymmetric distribution of membrane lipids in plasma membranes. Sphingomyelin, for example, is predominantly found in the outer leaflet, while phosphatidylethanolamine is enriched in the inner leaflet. This spatial organization is typically not preserved in artificial membranes such as GUVs used in reconstitution experiments of FGF2 membrane translocation (Steringer et al., 2017). In particular, the asymmetric distribution of PI(4,5)P_2_ in native plasma membranes is a striking difference compared to its symmetric distribution in artificial GUVs and, therefore, may contribute to the vastly different membrane translocation kinetics of FGF2 when these systems were compared. Hence, in the current study, we tested the hypothesis of PI(4,5)P_2_ transbilayer asymmetry to reduce the energy costs for FGF2-dependent membrane pore formation. The rationale for this was based on the concept of FGF2-dependent accumulation of the non-bilayer membrane lipid PI(4,5)P_2_ to produce a steep and spatially constrained electrochemical gradient across the plasma membrane that may facilitate the transformation of the stable lipid bilayer into a highly transient lipidic membrane pore (Gurtovenko and Vattulainen, 2007; Roesel et al., 2022). PI(4,5)P_2_-dependent FGF2 recruitment and oligomerization in cholesterol-rich domains has been shown to increase membrane tension (Lolicato et al., 2022). Based on our evidence-driven hypothesis that the FGF2 secretion machinery resides in liquid-ordered membrane domains (Lolicato and Nickel, 2022), this increase in membrane tension represents another parameter that is likely to facilitate pore opening (Akimov et al., 2017a; Akimov et al., 2017b). It is of note that a single FGF2 molecule is capable of recruiting up to 5 PI(4,5)P_2_ molecules through one high and several additional low affinity binding sites (Steringer et al., 2017). Since FGF2 is known to oligomerize in a membrane-dependent manner at least into tetramers (Steringer et al., 2017; Sachl et al., 2020; Singh et al., 2023), this would translate into the accumulation of a minimum of about 20 PI(4,5)P_2_ molecules in a spatially constrained manner. This in turn would produce a massive amount of local membrane stress in the vicinity of membrane-bound FGF2 oligomers at the inner plasma membrane leaflet based on (i) a steep electrochemical gradient of PI(4,5)P_2_ across the plasma membrane, (ii) a local accumulation of a lipid with non-bilayer characteristics and (iii) membrane tension based on high levels of cholesterol in regions of PI(4,5)P_2_-dependent FGF2 recruitment. We therefore hypothesized local and asymmetric accumulation of PI(4,5)P_2_ at FGF2 recruitment sites to lower the free energy costs required for the opening of the lipidic FGF2 membrane translocation pore.

Based on the above hypothesis, we developed an experimental in vitro system to produce model membranes that contain an asymmetric transbilayer distribution of PI(4,5)P_2_. Using an established assay to quantify FGF2-dependent membrane pore formation, we found an asymmetric distribution of PI(4,5)P_2_ to accelerate this process. Consistently, in cell-based experiments, we found FGF2 secretion to be inhibited under conditions compromising PI(4,5)P_2_ transbilayer asymmetry. Thus, our findings identify PI(4,5)P_2_ transbilayer asymmetry as an important parameter governing fast FGF2 membrane translocation kinetics as part of its unconventional mechanism of secretion from mammalian cells.

## Results

### Formation of artificial membranes with an asymmetric transbilayer distribution of PI(4,5)P_2_

To test the hypothesis of PI(4,5)P_2_ transbilayer asymmetry to play a role in FGF2 membrane translocation, we aimed at producing artificial membranes characterized by PI(4,5)P_2_ being present exclusively in the outer leaflet. We produced both LUVs and GUVs containing PI(4)P, a phosphoinositide to which FGF2 does not bind (Temmerman et al., 2008; Temmerman and Nickel, 2009). As illustrated in Fig. 1A, the approach was to convert PI(4)P into PI(4,5)P_2_ by an enzymatic reaction catalyzed by the lipid kinase PIP5K1C (Ishihara et al., 1998). In the presence of Mg^2+^ ions and ATP, assuming exclusion of the enzyme from the lumina of LUVs and GUVs, we aimed at the production of PI(4,5)P_2_ exclusively in the outer leaflet of LUVs and GUVs. Two types of analyses were conducted to test successful production of PI(4,5)P_2_: FGF2-GFP binding to LUVs analyzed by a previously established assay using flow cytometry (Fig. 1B) and FGF2-GFP binding to LUVs analyzed by biochemical sedimentation assays (Fig. 1C). As demonstrated in Fig. 1B, in comparison to PI(4,5)P_2_, FGF2-GFP does not bind to PI(4)P. Following the addition of Mg^2+^ ions as part of the enzymatic conversion of PI(4)P into PI(4,5)P_2_, relative binding efficiencies of FGF2-GFP were generally lower due to clustering of liposomes having an impact on quantification using flow cytometry. Nevertheless, following conversion of PI(4)P into PI(4,5)P_2_, similar levels of FGF2-GFP binding were observed compared to LUVs to which PI(4,5)P_2_ had been added during their formation. By contrast, under mock conditions, FGF2-GFP failed to bind to liposomes that were treated with ATP and Mg^2+^ ions in the absence of PIP5K1C. Similar results were obtained when biochemical sedimentation experiments were done to quantify FGF2-GFP binding to LUVs (Fig.1C). With this read-out, FGF2-GFP binding efficiencies were similar under standard conditions and in the presence of ATP, and Mg^2+^ ions and PIP5K1C. Upon enzymatic conversion of PI(4)P into PI(4,5)P_2_, efficient FGF2-GFP binding could be observed that was similar to LUVs to which PI(4,5)P_2_ had been added during their formation. Finally, we extended this system to GUVs analyzed by confocal microscopy (Fig. 1D and 1E). While FGF2-GFP did bind efficiently to PI(4,5)P_2_-containing GUVs, it bound to GUVs made with PI(4)P only after enzymatic conversion into PI(4,5)P_2_. The combined experiments shown in Fig. 1 demonstrate the feasibility of the enzymatic conversion approach to generate PI(4,5)P_2_ from PI(4)P in the outer leaflet of both LUVs and GUVs.

**Figure 1.**
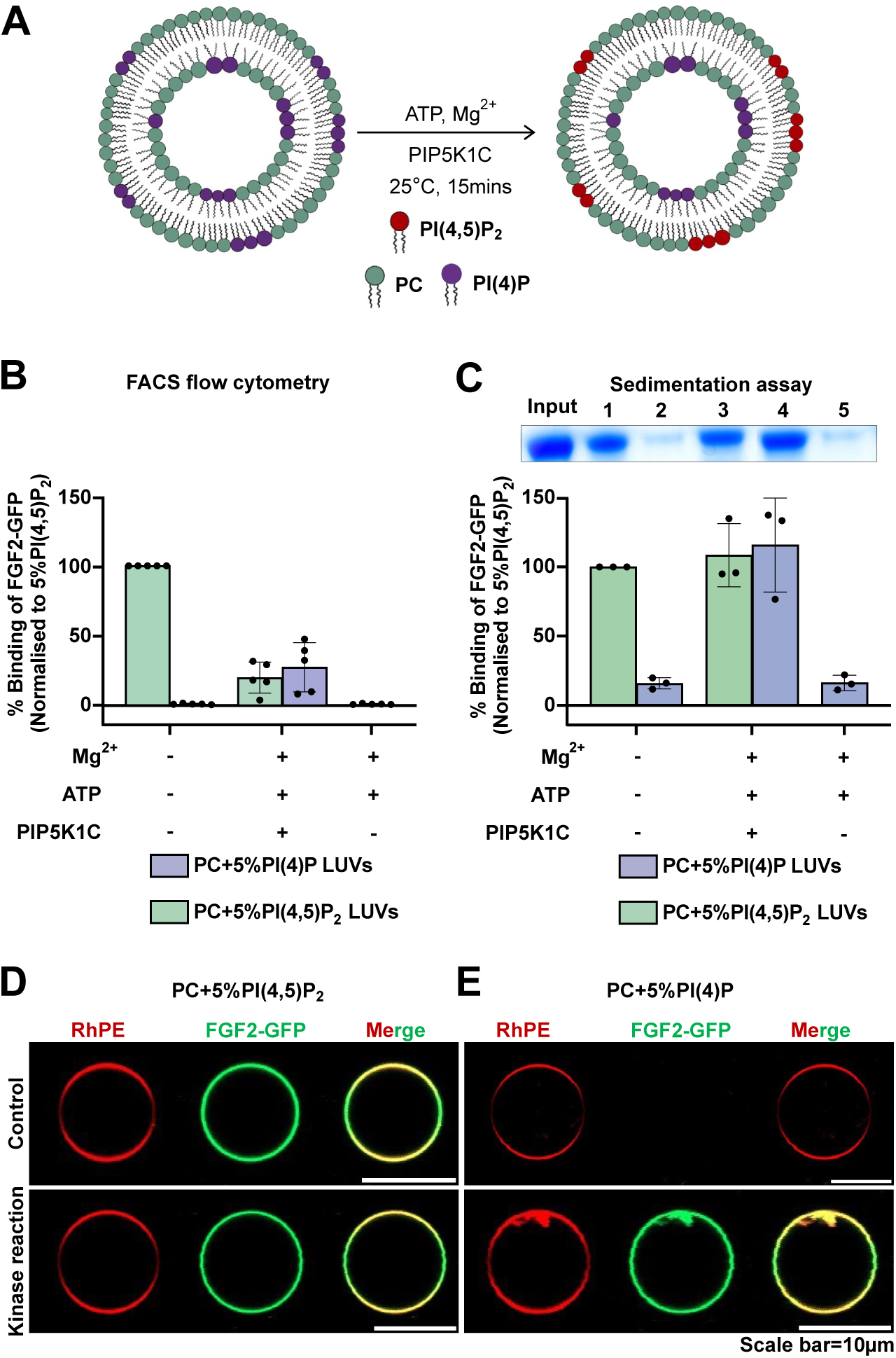
Enzymatic approach to generate PI(4,5)P₂ asymmetric vesicles (A) Schematic representation of enzymatic approach to generates transbilayer PI(4,5)P₂ asymmetry. Reaction utilizes PI(4)P containing liposomes with Mg²^+^, ATP, and PIP5K1C to selectively phosphorylate PI(4)P to PI(4,5)P₂ on the outer leaflet of the vesicles. Large unilamellar vesicles (LUVs) were prepared with 5 mol% PI(4,5)P₂ or 5 mol% PI(4)P in PC background. LUVs with 5 mol% PI(4,5)P₂ were subjected to kinase reaction or left untreated. Similarly, LUVs with 5 mol% PI(4)P were subjected to kinase reaction or mock reaction (without PIP5K1C) or left untreated. After the reaction, vesicles were incubated with FGF2-GFP for 60 minutes, and unbound protein was removed by centrifugation before analysis. (B) Analysis of FGF2-GFP binding using FACS flow cytometry. Each dot represents a biological replicate with 2–3 technical replicates per condition. For each experiment, average signal from 5 mol% PI(4,5)P₂ was set to 100%, and all values were normalized accordingly. Mean values with standard deviation (S.D.) are presented. (C) Analysis of FGF2-GFP binding using Sedimentation assay. The bound fraction of FGF2-GFP was analysed by SDS-PAGE and detected with Coomassie staining. Gel was imaged by EPSON 4870 scanner. Data were normalized to 5 mol% PI(4,5)P₂ within each experiment. Mean values with S.D. are presented. (D) Giant unilamellar vesicles (GUVs) were prepared with 5 mol% PI(4,5)P₂ or (E) 5 mol% PI(4)P along with membrane marker Rhodamine-PE, in PC background. GUVs were subjected to kinase reaction or left untreated. After the kinase reaction, vesicles were incubated with FGF2-GFP for 1h and imaged with confocal microscope to access the conversion of PI(4)P to PI(4,5)P₂. Representative confocal images of FGF2-GFP binding to GUVs with and without kinase reaction are shown.

### Analysis of the transbilayer distribution of PI(4,5)P_2_ in large unilamellar vesicles

To validate the approach of generating artificial membranes with an asymmetric transbilayer distribution of PI(4,5)P_2_, we established a quantitative procedure that was based on sequential disintegration and reformation of LUVs followed by FGF2-GFP binding experiments (Fig. 2). LUVs were disintegrated using 40% methanol in chloroform. Lipids were extracted from the organic phase and used to reform liposomes followed by the quantitative assessment of FGF2 binding to PI(4,5)P_2_. As illustrated in Fig. 2A, we expected a reduction of surface PI(4,5)P_2_ in asymmetric LUVs due to dilution of PI(4,5)P_2_ into the inner leaflet (Fig. 2A, lower panel). By contrast, a disintegration and reformation procedure would not alter the surface amounts of PI(4,5)P_2_ in LUVs that were formed with a symmetric distribution of PI(4,5)P_2_ (Fig. 2A, upper panel). As shown in Fig. 2B with non-treated LUVs shown in grey and disintegrated/reformed LUVs in light red, the normal binding specificities of FGF2-GFP towards PI(4,5)P_2_ versus PI(4)P were observed (Fig. 2B; conditions a versus b) based on flow cytometry (Temmerman et al., 2008; Temmerman and Nickel, 2009). LUVs containing PI(4,5)P_2_, treated with PIP5K1C, Mg^2+^ and ATP, when measured without disintegration/reformation showed lower binding efficiencies towards FGF2-GFP (Fig. 2B, condition c, grey bar). Following disintegration/reformation, they went up to normal levels (Fig. 2B, condition c, light red bar). This is because disintegration/reformation procedure apparently removed Mg^2+^ ions, as a result of which, binding intensity recovered to normal when compared to non-treated PI(4,5)P_2_ vesicle (Fig. 2B, condition a, grey bar versus c, light red bar). For LUVs containing PI(4)P, treated with PIP5K1C, Mg^2+^ and ATP, when directly analyzed for FGF2-GFP binding efficiencies, similar levels were observed compared to LUVs containing PI(4,5)P_2_ (Fig. 2B, condition d, grey bar). Most importantly, when LUVs containing PI(4)P, treated with PIP5K1C, Mg^2+^ and ATP were subjected to the disintegration/reformation procedure, FGF2-GFP binding efficiencies did not went up as observed for preformed PI(4,5)P_2_ LUVs but rather substantially down (Fig. 2B, condition d, light red bar). A comparison of FGF2 binding efficiencies with LUVs containing different concentrations of PI(4,5)P_2_ (Fig. 2C) indicated that such LUVs had a surface concentration of enzymatically produced PI(4,5)P_2_ between 2 and 3 mol%. Compared to the starting condition of 5 mol%, this indicated that the disintegration/reformation procedure indeed caused dilution of PI(4,5)P_2_ into the inner leaflet. This validated our approach with LUVs formed with PI(4)P followed by enzymatic conversion into PI(4,5)P_2_ being characterized by an asymmetric transbilayer distribution of PI(4,5)P_2_ with at least the vast majority of this membrane lipid being present in the outer leaflet of LUVs.

**Figure 2.**
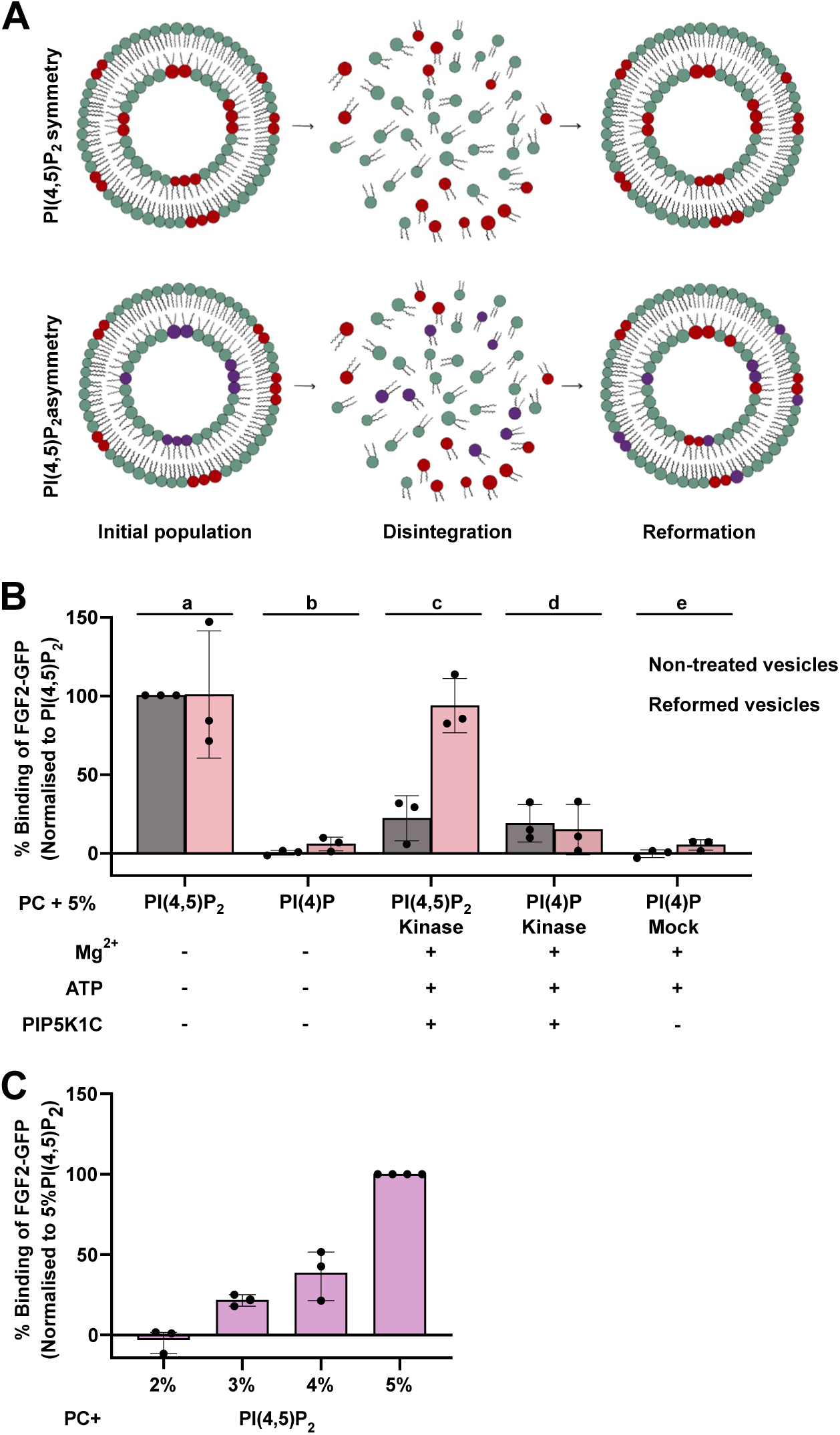
Testing transbilayer PI(4,5)P_2_ asymmetry in large unilamellar vesicles (A) Schematic representation of asymmetry test using vesicle disintegration/reformation method. Vesicles were disintegrated with 40% MeOH in CHCl₃ and then reconstructed to assess their binding towards FGF2-GFP. (B) Quantification of FGF2-GFP binding to PC LUVs using FACS flow cytometry. Vesicles were incubated with FGF2-GFP for 30 minutes, and unbound protein was removed by centrifugation before analysis. Binding was assessed for LUVs with 5 mol% PI(4,5)P₂ (sub-panel a), 5 mol% PI(4)P (sub-panel b), 5 mol% PI(4,5)P₂+Mg^2+^+ATP+PIP5K1C (sub-panel c), 5 mol% PI(4)P+Mg^2+^+ATP+PIP5K1C (sub-panel d), 5 mol% PI(4)P+Mg^2+^+ATP (sub-panel e). Gray bars represent initial/non -treated population of LUVs and light red bar represent reformed LUVs. Each dot represents a biological replicate with 2 technical replicates per condition. For each experiment, the average signal from non-treated/initial 5 mol% PI(4,5)P₂ LUV population was set to 100%, and all values were normalized accordingly. Mean values with standard deviation (S.D.) are presented. (C) Analysis of FGF2-GFP binding to PC LUVs with increasing mol% PI(4,5)P_2_ using FACS flow cytometry. Vesicles were incubated with FGF2-GFP for 30 minutes, and unbound protein was removed by centrifugation before analysis. Each dot represents a biological replicate with 2–3 technical replicates per condition. For each experiment, average signal from 5 mol% PI(4,5)P₂ was set to 100%, and all values were normalized accordingly. Mean values with S.D. are presented.

### PIP5K1C mediated conversion of PI(4)P into PI(4,5)P_2_ on the membrane surface of both LUVs and GUVs with a plasma membrane-like lipid composition

The basic approach of converting PI(4)P at the outer leaflet of LUVs and GUVs into PI(4,5)P_2_ by the enzymatic action of PIP5K1C to produce artificial membranes with an asymmetric transbilayer distribution of PI(4,5)P_2_ was established with a simple bulk lipid composition consisting of phosphatidylcholine (Figs. 1 and 2). To prepare for functional experiments such as FGF2-dependent membrane pore formation, we aimed at making the lipid composition more complex including cholesterol and sphingomyelin mimicking plasma membranes, the subcellular site of FGF2 membrane translocation. On the one hand, we produced LUVs with a plasma membrane like lipid composition containing either PI(4)P or PI(4,5)P_2_ (Fig. 3A). To quantify lipid binding, we used the FGF2-Halo construct instead of GFP-tagged FGF2, as it has been shown to exhibit minimal liposome tethering in LUVs with complex lipid compositions (Lolicato et al., 2022). When FGF2-Halo binding studies were conducted based upon biochemical sedimentation experiments, the binding efficiency of FGF2-Halo towards PI(4,5)P_2_ containing LUVs (light green bars) was set as a reference point (Fig. 3A, light green bar, condition 1). A substantial reduction in FGF2-Halo binding efficiency was observed for LUVs with a plasma membrane like lipid composition containing PI(4)P (Fig. 3A, light pink bars). As shown in Fig. 3A (condition 3 and 4), a binding efficiency of FGF2-Halo comparable to PI(4,5)P_2_ LUVs could be recovered when PI(4)P containing LUVs were incubated with PIP5K1C, Mg^2+^ and ATP, converting PI(4)P into PI(4,5)P_2_. Similar results were obtained with GUVs containing a plasma membrane like lipid composition supplemented with either PI(4,5)P_2_ or PI(4)P (Fig. 3B and 3C). While FGF2-GFP did bind efficiently to GUVs containing PI(4,5)P_2_ (Fig. 3B, control), absence of binding was observed when GUVs contained PI(4)P (Fig. 3C, control). Of note, conversion of PI(4)P into PI(4,5)P_2_ restored FGF2-GFP binding to GUVs (Fig. 3B and 3C, kinase reaction). A similar trend was observed when FGF2-Halo was used in GUV experiments. The combined experiments shown in Figs. 1, 2 and 3 established that PI(4)P into PI(4,5)P_2_ conversion mediated by PIP5K1C works both in a simple PC background (Figs. 1 and 2) and in a more complex membrane lipid background mimicking the plasma membrane lipid composition (Fig. 3). In addition, incubations of PIP5K1C in the presence of ATP and Mg^2+^ ions did not compromise membrane integrity of GUVs throughout the experiment as indicated by the exclusion of a small fluorescent tracer, Alexa-647 (Fig. 3D and 3E as well as videos 1 and 2). Similar to the experiments shown in Fig. 1 and 2, this indicates that PIP5K1C acted on PI(4)P exclusively from the outside, producing PI(4,5)P_2_ in the outer leaflet of GUVs only. Hence, alike the experiments shown in Figs 1 and 2, this procedure produced GUVs with an asymmetric transbilayer distribution of PI(4,5)P_2_.

**Figure 3:**
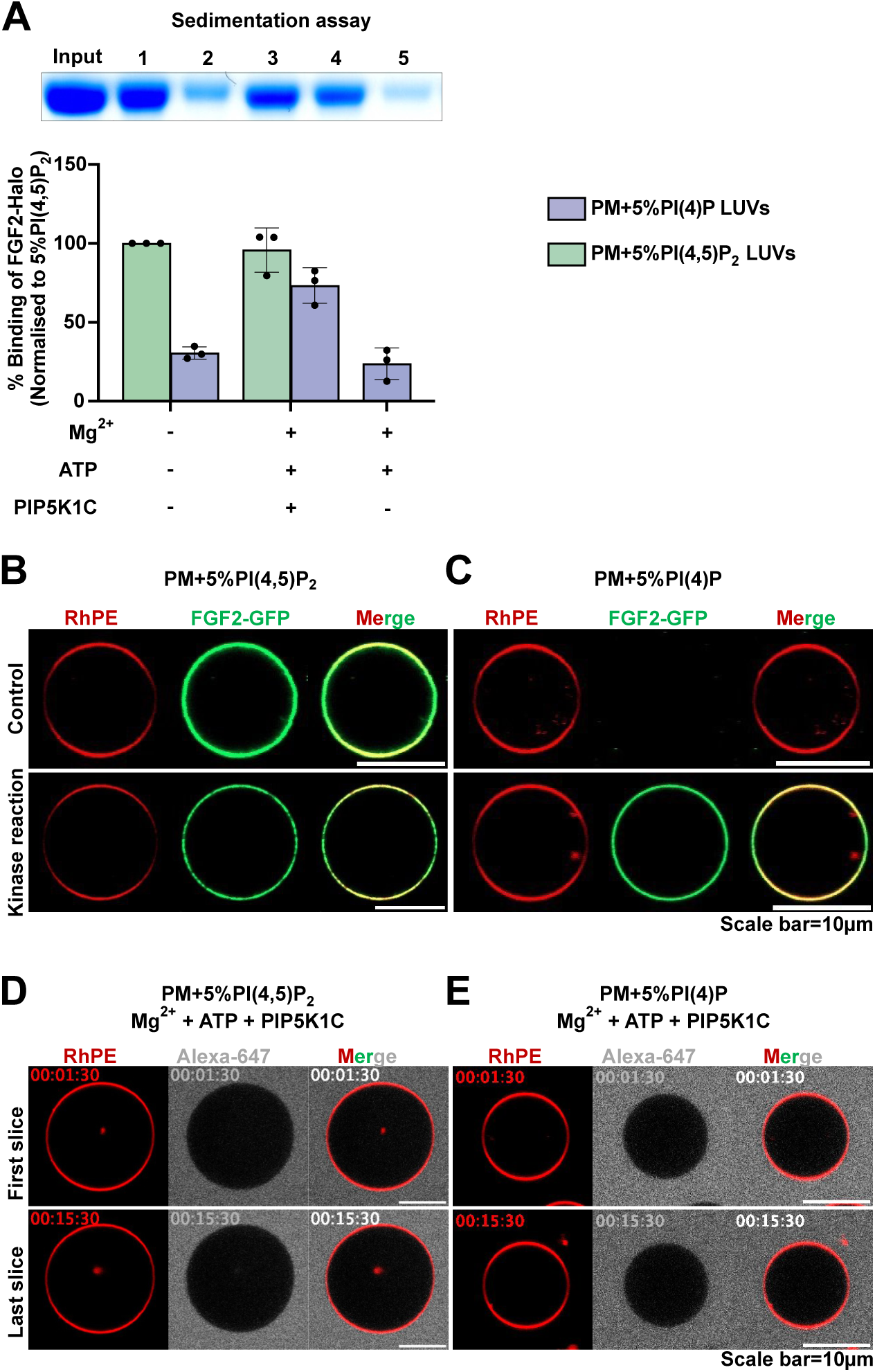
Generation of PI(4,5)P₂ asymmetric vesicles with plasma membrane like lipid composition (A) Large unilamellar vesicles (LUVs) were prepared with 5 mol% PI(4,5)P₂ or 5 mol% PI(4)P in plasma membrane (PM) like background. These LUVs were subjected to kinase reaction or mock reaction (without PIP5K1C) or left untreated. After the reaction, vesicles were incubated with FGF2-Halo for 3 hours, and unbound protein was removed by centrifugation before analysis using a sedimentation assay. The bound fraction of FGF2-Halo was analyzed by SDS-PAGE and detected with Coomassie staining. The gel was imaged using EPSON 4870 scanner. Data were normalized to 5 mol% PI(4,5)P₂ within each experiment. Mean values with standard deviation are presented. (B) Giant unilamellar vesicles (GUVs) were prepared with 5 mol% PI(4,5)P₂ or (C) 5 mol% PI(4)P along with membrane marker Rhodamine-PE, in PM like background. GUVs were subjected to kinase reaction or left untreated. After the kinase reaction, vesicles were incubated with FGF2-GFP for 1h and imaged with confocal microscope to access the enzymatic conversion of PI(4)P to PI(4,5)P₂. Representative confocal images of FGF2-GFP binding to GUVs with and without kinase reaction are shown. (D) To test the asymmetry in GUVs, vesicles with 5 mol% PI(4,5)P₂ or (E) 5 mol% PI(4)P along with membrane marker Rhodamine-PE, in PM like background, were immobilized on glass bottom ibidi chambers. Vesicles were monitored after the addition of Mg^2+^ + ATP + PIP5K1C and small tracer dye Alexa-647 in real time for 15 mins using confocal microscope. Each time frame is separated by 1min time. First slice was recorded at 1.5 min and last time slice was recoded at 15 mins. Representative confocal images of GUVs with kinase reaction remains intact during the entire course of kinase reaction.

### PI(4,5)P_2_ asymmetry facilitates FGF2-dependent membrane pore formation in reconstitution experiments

To test the hypothesis as to whether the kinetics of FGF2 membrane translocation are affected by the transbilayer distribution of PI(4,5)P_2_, we used the GUV system with a plasma membrane like lipid composition (Fig. 3) in combination with imaging assays detecting FGF2-induced membrane pore formation established previously (Steringer et al., 2012; Steringer et al., 2017; Lolicato et al., 2024). Three experimental conditions were compared (i) GUVs with a symmetric transbilayer distribution, (ii) GUVs with an asymmetric transbilayer distribution of PI(4,5)P_2_ and (iii) GUVs with a symmetric transbilayer distribution treated with PIP5K1C, ATP and Mg^2+^ ions as a control (Fig. 4). In Fig. 4A, examples of still images of GUVs are shown at an early time point (20 to 25 min) of incubation with FGF2-GFP. Along with a fluorescently labeled membrane lipid (RhPE), FGF2-GFP binding to the membrane surfaces of GUVs was visible for all three conditions. Importantly, at these time points, membrane pore formation was not detectable as indicated by the luminal exclusion of the small fluorescent tracer, Alexa-647. In Fig. 4B, examples are given for individual GUVs from later time points along with the incubation time that led to FGF2-dependent membrane pore formation, indicated by luminal penetration of the small fluorescent tracer molecule (Alexa-647). For the given examples, this event occurred after about 2 hours for the GUV with a symmetric transbilayer distribution, after 40 min for the GUV with an asymmetric transbilayer distribution of PI(4,5)P_2_ and after 1 hour and 27 min for the GUV with a symmetric transbilayer distribution treated with PIP5K1C, ATP and Mg^2+^ ions as a control (Fig. 4B). The videos 3, 4 and 5 cover the incubation time for all three of these conditions, visualizing the dynamics of this process with pore formation at different time points depending on the experimental conditions described above.

**Figure 4.**
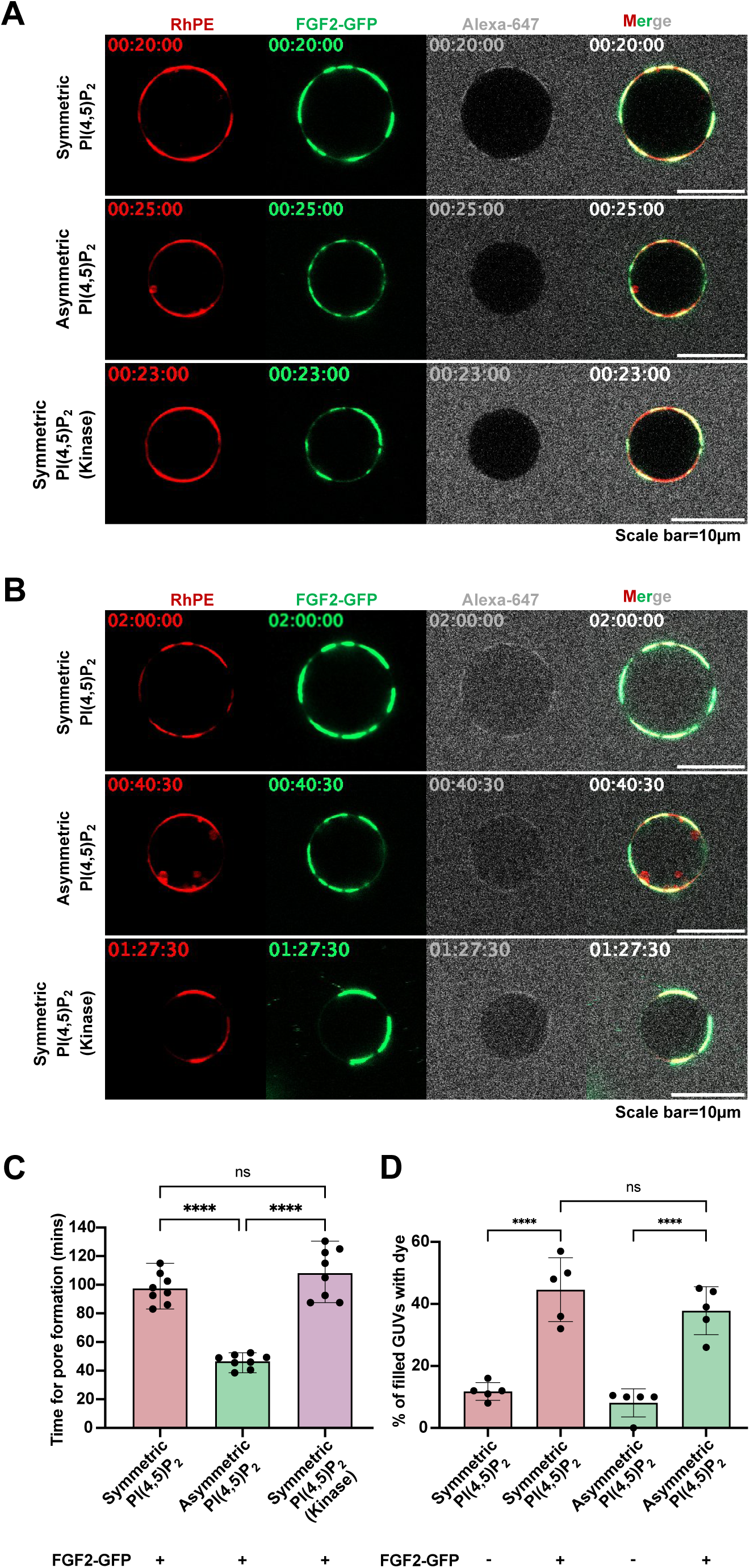
FGF2 translocation kinetics depend on the asymmetric distribution of PI(4,5)P₂ in vesicles Giant unilamellar vesicles (GUVs) were prepared with 5 mol% PI(4,5)P₂ or 5 mol% PI(4)P along with membrane marker Rhodamine-PE, in PM like background. Small trace dye Alexa-647 was added to record the event of pore formation for kinetic measurement. Three conditions were compared for FGF2 translocation kinetic study; GUVs with symmetric PI(4,5)P₂, asymmetric PI(4,5)P₂ and symmetric PI(4,5)P₂ with Mg^2+^ + ATP + PIP5K1C as a control. After addition of FGF2-GFP, time was marked as 0 min and vesicle was given 10-20 mins to immobilize and imaging was started at 20-25 mins till 01:30-02:00 h. (A) First recorded time frame for PM vesicles with symmetric or asymmetric PI(4,5)P_2_ or symmetric vesicles with reaction ingredients. (B) Still frame at the real time of pore formation for PM vesicles with symmetric or asymmetric PI(4,5)P_2_ or symmetric vesicles with reaction ingredients. (C) Quantification and statistical analysis of pore formation kinetics for FGF2-GFP in symmetric, asymmetric PI(4,5)P_2_ and symmetric vesicles with reaction ingredients. Each dot represents a single GUV pore formation in real time. (D) Quantification and statistical analysis of FGF2-GFP dependent pore formation. Each dot represents percentage of GUVs with membrane pores with a ratio of luminal versus exterior Alexa647 tracer ≥0.6. In each independent experiment 20-59 GUVs were examined. Mean values with S.D. are presented. For statistical test, ordinary one-way ANOVA with Turkey test was performed in Prism. Not significant (ns) P > 0.5, **** P ≤ 0.0001. Data distribution was assumed to be normal.

To test the significance of the observed differences depicted in Fig. 4A and 4B, we recorded the time intervals of membrane pore formation for 8 GUVs per condition, derived from 4–7 independent experiments. These measurements were conducted for each of the three experimental conditions described (Fig. 4C). This analysis revealed a statistically significant shortening of time intervals required for FGF2-dependent membrane pore formation when GUVs with an asymmetric transbilayer distribution of PI(4,5)P_2_ (mean value of about 45 min) were compared with those having a symmetric distribution of membrane lipids (mean value of about 95 min). By contrast, the time intervals for FGF2-dependent membrane pore formation measured for GUVs with a symmetric transbilayer distribution and those additionally treated with PIP5K1C, ATP and Mg^2+^ ions did not differ with a mean value of about 105 min for the latter (Fig. 4C). Of note, the total efficiency of FGF2-dependent membrane pore formation observed as the percentage of GUVs containing the Alexa-647 tracer at the end of the measurements did not differ between GUVs with a symmetric versus those having an asymmetric distribution of PI(4,5)P_2_ (Fig. 4D). These findings suggest transbilayer asymmetry of PI(4,5)P_2_ to represent an important parameter for the thermodynamic properties of this system, governing the kinetics of FGF2-induced membrane pore formation.

### Disturbing plasma membrane transbilayer asymmetry of PI(4,5)P_2_ in intact cells

To challenge the results from the biochemical reconstitution experiments shown in Figs. 1-4 and the accompanying videos 3-5, we went further to test as to whether plasma membrane transbilayer asymmetry of PI(4,5)P_2_ in intact cells plays a role in unconventional secretion of FGF2. We established a procedure to disturb the asymmetric distribution of PI(4,5)P_2_ in intact cells that was inspired by the delivery of functionalized phosphoinositides to cells (Golebiewska et al., 2008; Schultz, 2023; Schultz and Brügger, 2023). As illustrated in Fig. 5A, PI(4,5)P_2_ and Phosphatidylserine (PS) as a control were added to cells in an aqueous solution most likely forming micelles (PI(4,5)P_2_) or liposomal vesicles (PS). We reasoned cellular delivery would occur at the outer plasma membrane leaflet, compromising transbilayer asymmetry of both endogenous membrane lipids that under normal conditions localize exclusively to the inner plasma membrane leaflet in living cells. Delivery was experimentally confirmed by confocal imaging using a recombinant PH domain based fluorescent sensor for PI(4,5)P_2_ (PHx2-Halo; (Lolicato et al., 2022)) and a fluorescent variant of Annexin V for PS. As shown in Fig. 5B, as opposed to mock conditions, both PI(4,5)P_2_ (left-hand panels) and PS (right-hand panels) were readily detectable by the recombinant fluorescent sensors that were added to cells from outside, detecting these lipids in the outer plasma membrane leaflet. Based on our calculations of PI(4,5)P₂ levels for individual cells (see Materials and Methods), we added a three-fold excess of PI(4,5)P₂. We reasoned that this concentration would effectively disrupt PI(4,5)P₂ transbilayer asymmetry, which is consistent with the signal observed using the PHX2-Halo protein.

**Figure 5.**
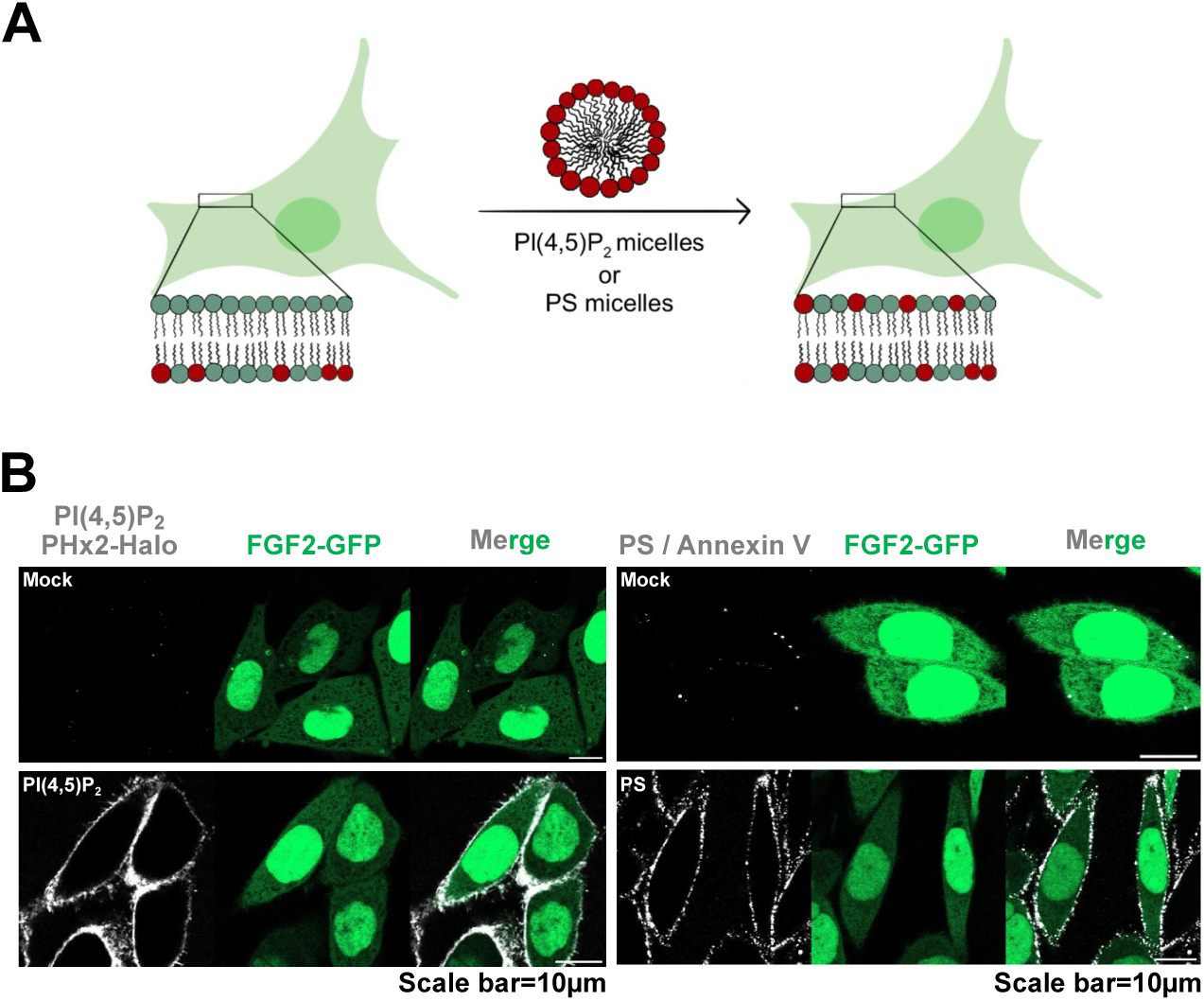
Disruption of cellular transbilayer asymmetry of PI(4,5)P₂ or PS (A) Schematic representation of the disruption of plasma membrane PI(4,5)P_2_ or PS transbilayer asymmetry using PI(4,5)P_2_ or PS micelles/vesicles. (B) Representative confocal images of CHO-K1 cells expressing FGF2-GFP in doxycycline-dependent manner, with disrupted PI(4,5)P_2_ or PS asymmetry. On left of panel B, are fixed cells stained with high affinity PI(4,5)P_2_ binding protein PHX2-Halo (40nM). The Halo tag was visualized using Halo ligand Alexa 660 (75nM). Cells were fixed using 3% PFA in PBS for 13mins at room temperature. PI(4,5)P_2_ signal was observed only upon addition of PI(4,5)P_2_ micelles, confirming the successful disruption of PI(4,5)P_2_ asymmetry. On right of panel B are live cells stained with PS binding protein AnnexinV-Alexa647 in presence of 2mM Ca^2+^ in Live cell imaging solution. PS signal was observed only upon addition of PS micelles, confirming the successful disruption of PS asymmetry.

### A quantitative assay measuring an acute wave of FGF2 secretion from cells

We have previously established a number of quantitative assays measuring FGF2 transport into the extracellular space based upon, for example, cell surface biotinylation as well as confocal and TIRF microscopy (Seelenmeyer et al., 2005; Zehe et al., 2006; Temmerman et al., 2008; Torrado et al., 2009; Ebert et al., 2010; Steringer et al., 2012; Müller et al., 2015; Zacherl et al., 2015; La Venuta et al., 2016; Dimou et al., 2019). These experimental systems were designed to quantify FGF2 on cell surfaces at equilibrium, with the experiment initiated by doxycycline-dependent induction of FGF2 expression and cell surface FGF2 quantified after the system reached equilibrium. In the context of the current study, we aimed at developing a novel experimental system based upon a single cell analysis using confocal microscopy, quantifying an acute wave of FGF2 translocation to cell surfaces within a short time interval in the range of minutes (Fig. 6). Cells were treated for 16 hours with doxycycline to induce FGF2-GFP expression with an equilibrium distribution between the cytoplasm, the nucleus and the cell surface. While the first two subcellular localizations were directly observable by GFP fluorescence, cell surface FGF2-GFP was detected on intact cells using fluorescently labeled anti-GFP antibodies (Fig. 6A). In a second step, cells were treated with heparin molecules with a high affinity towards FGF2 to wash away and remove FGF2-GFP from cell surfaces (Fig. 6B). Following the heparin wash, time was marked as 0 min. Subsequently, a new wave of FGF2-GFP transported to cell surfaces could be observed as shown in Fig. 6C. Reappearance of FGF2-GFP on cell surfaces occurred within 30 min of incubation. This setup worked efficiently with FGF2-GFP removed quantitatively from cell surfaces by a stringent heparin wash (Fig. 6A versus 6B) and full recovery of the cell surface population of FGF2-GFP within 30 min of incubation (Fig. 6C), the latter representing an acute wave of FGF2 secretion into the extracellular space.

**Figure 6.**
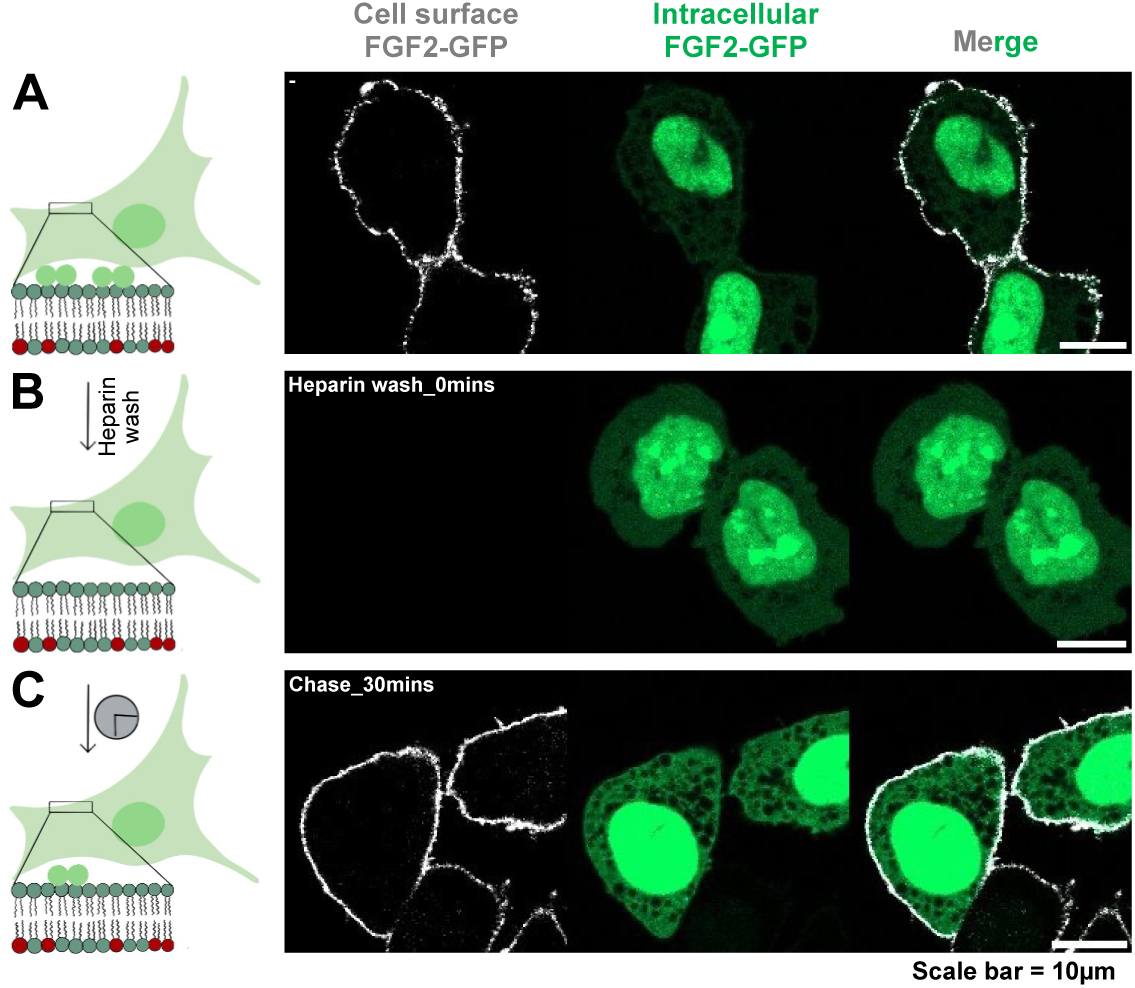
A quantitative assay measuring an acute wave of FGF2 secretion from cells Schematic representation of assay used to measure FGF2 secretion from CHO-K1 cells. CHO-K1 cells were induced to express FGF2-GFP overnight by adding 1.1:1000 doxycycline. (A) FGF2-GFP localization in equilibrium, with FGF2-GFP distributed in the nucleus, cytosol, inner and outer leaflet of plasma membrane. On the right side there is a representative confocal image of fixed cells after overnight doxycycline induction. Surface FGF2-GFP was detected using rabbit polyclonal anti-GFP-Alexa647 antibody at 1:400 dilution. (B) Surface FGF2-GFP was removed using 1 mg/mL heparin. On the right side there is a representative confocal image of fixed cells at 0 min after heparin wash. (C) Recovery of FGF2-GFP on the cell surface was monitored over time post heparin wash. On the right side there is a representative confocal image of fixed cells at 30 mins after heparin wash. Cells were fixed using 3% PFA in PBS for 13 mins at room temperature. Using this assay, surface FGF2-GFP can be monitored at the single cell resolution.

### Transbilayer asymmetry of PI(4,5)P_2_ in plasma membranes is required for FGF2 secretion from cells

Using the experimental setup described in Fig. 6, we tested as to whether disturbance of the asymmetric distribution of PI(4,5)P_2_ in the plasma membranes of cells directly affects the efficiency of FGF2 membrane translocation into the extracellular space (Fig. 7). Four conditions were compared to quantify FGF2 secretion within a 30 min time interval with (i) untreated cells, (ii) mock-treated cells, (iii) cells treated with exogenous PI(4,5)P_2_ and (iv) cells treated with exogenous PS, using the procedures established in Fig. 5. In Fig. 7A, representative example images for each of the four experimental conditions are shown. For untreated and mock-treated as well as PS-treated cells, efficient reappearance of FGF2-GFP on cell surfaces could be observed, demonstrating normal FGF2-GFP transport efficiencies into the extracellular space under these conditions. By contrast, in cells treated with exogenous PI(4,5)P_2_ to disturb transbilayer asymmetry, FGF2-GFP failed to reappear at the cell surface, demonstrating FGF2 membrane translocation to be blocked under this experimental condition. Quantification along with statistics comparing the four experimental conditions mentioned above were done with single cell resolution (Fig. 7B). Every single data point corresponds to the quantitative analysis of cell surface FGF2-GFP in individual cells. At time point 0 min (Fig. 7B, left-hand side), FGF2-GFP was hardly detectable on cell surfaces, demonstrating the effectiveness of the heparin wash, marking the start condition of the experiment. After 30 min of incubation (Fig. 7B, right-hand side), substantial amounts of FGF2-GFP on cell surfaces could be detected for untreated, mock-treated and PS-treated cells without any statistically significant differences. By contrast, cells treated with exogenous PI(4,5)P_2_ were characterized by very low levels of FGF2-GFP on cell surfaces similar to the one detected at 0 min of incubation. The observed differences to mock-and PS-treated cells were highly statistically significant, demonstrating the strong specificity of the effect exerted by PI(4,5)P_2_. Thus, the transbilayer asymmetry of PI(4,5)P_2_ in plasma membranes is a key parameter enabling efficient FGF2 membrane translocation into the extracellular space.

**Figure 7.**
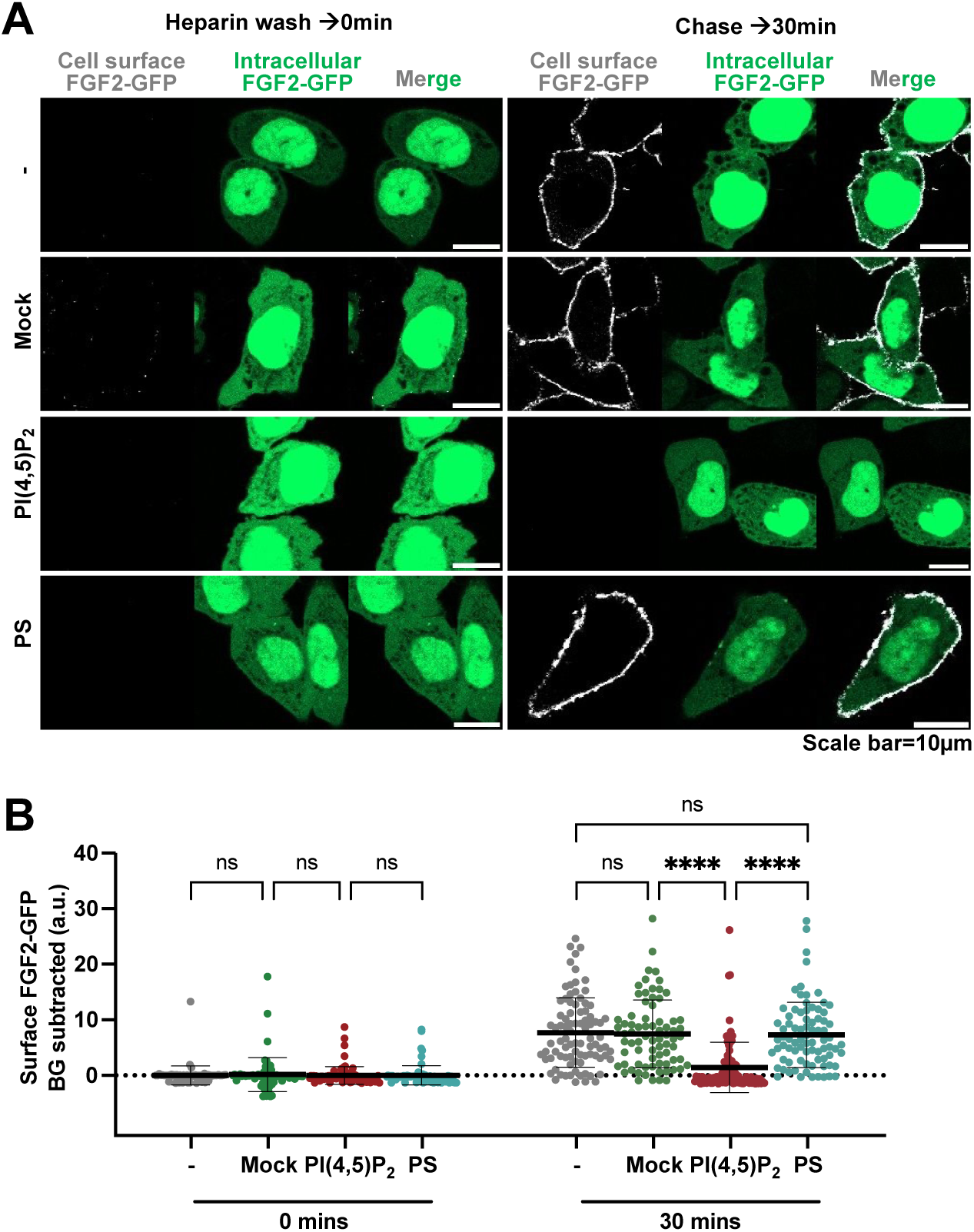
Disruption of transbilayer PI(4,5)P₂ asymmetry inhibits FGF2-GFP secretion in cells CHO-K1 cells were induced to express FGF2-GFP overnight by adding 1.1:1000 doxycycline. (A) Representative confocal images of non-treated, mock-treated, PI(4,5)P_2_-treated, and PS-treated cells. Cells were treated with 1 mg/mL heparin to remove surface FGF2-GFP. On left side of the panel are the representative confocal images of fixed cells at 0 min after the heparin wash. On the right side of the panel there are confocal images of the fixed cells at 30 mins after the heparin wash. Cells were fixed using 3%PFA in PBS for 13 mins at room temperature. Surface FGF2-GFP was detected using rabbit polyclonal anti-GFP-Alexa647 antibody at 1:400 dilution. (B) Quantification and statistical analysis of surface FGF2-GFP levels for individual cells that were left un-treated, mock-treated, PI(4,5)P_2_-treated, and PS-treated. Each dot represents a single cell coming from 3 independent experiments. Unspecific background from antiGFP-AlexFluor-647 signal after heparin wash for each condition is subtracted. Mean values with standard deviation are presented. For statistical test, One-way ANOVA (Brown-Forsythe and Welch ANOVA test) with Dunnett T3 (n>50 per group) test was performed in Prism. Not significant (ns) P > 0.5, **** P ≤ 0.0001. Data distribution was assumed to be normal.

To further challenge these findings, we analyzed the lifetime of exogenously added PI(4,5)P_2_ on cell surfaces in the course of the experiment, using the procedures described in Fig. 5. As shown in Fig. 8A, starting at about 40 mins following addition of PI(4,5)P_2_, the cell surface levels of PI(4,5)P_2_ started to decrease and were hardly detectable at 60 min of incubation. This suggested that, at time points later than our reference point of 30 mins, cells were capable of re-establishing PI(4,5)P_2_ asymmetry in their plasma membranes, presumably by endocytosis of exogenously added PI(4,5)P_2_. Intriguingly, as shown in Fig. 8B, FGF2-GFP translocation to cell surfaces recovered when exogenously added PI(4,5)P_2_ disappeared from cell surfaces at time points of 40 to 60 mins. These findings corroborate our conclusion that the failure of FGF2-GFP to reach cell surfaces at 30 min of incubation is due to the loss of PI(4,5)P_2_ asymmetry, exerted by the addition of exogenous PI(4,5)P_2_ from outside cells.

**Figure 8.**
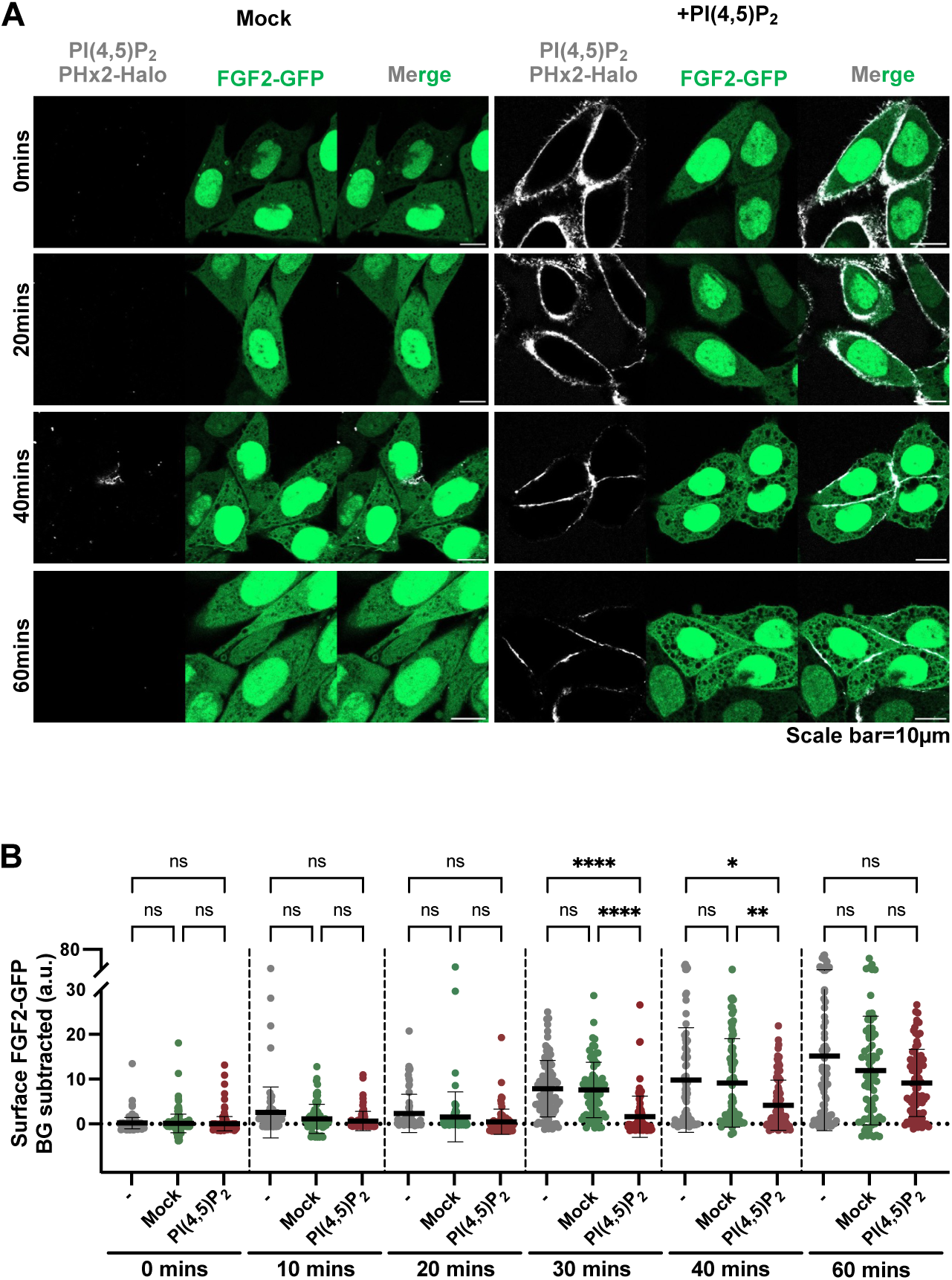
FGF2-GFP secretion is modulated by PI(4,5)P₂ levels at the plasma membrane surface CHO-K1 cells were induced to express FGF2-GFP overnight by adding 1.1:1000 doxycycline. (A) Representative confocal images of mock-treated and PI(4,5)P_2_-treated CHO-K1 cells. Cells were fixed either at 0 mins, 20 mins, 40 mins and 60 mins after disruption of plasma membrane PI(4,5)P_2_ asymmetry. Fixed cells were stained with high affinity PI(4,5)P_2_ binding protein PHX2-Halo (40 nM). The Halo tag was visualized using Halo ligand Alexa 660 (75 nM). (B) Quantification and statistical analysis of surface FGF2-GFP levels for individual cells that were left un-treated, mock-treated, and PI(4,5)P_2_-treated. Cells were fixed at 0 mins, 20 mins, 30 mins, 40 mins, and 60 mins after the heparin wash. Cells were fixed using 3%PFA in PBS for 13 mins at room temperature. Surface FGF2-GFP was detected using rabbit polyclonal anti-GFP-Alexa647 antibody at 1:400 dilution. Each dot represents a single cell coming from 3-9 independent experiments. Unspecific background from antiGFP-AlexaFluor-647 signal after heparin wash for each condition was subtracted. Mean values with standard deviation are presented. For statistical test, One-way ANOVA (Brown-Forsythe and Welch ANOVA test) with Dunnett T3 (n>50 per group) test was performed in Prism. Not significant (ns) P > 0.5, **** P ≤ 0.0001. Data distribution was assumed to be normal.

## Discussion

A general hallmark of biological membranes from a broad range of organisms is their structural organization in laterally partitioned nanodomains as well as their transbilayer asymmetry of membrane lipid distribution between the two leaflets (Bretscher, 1972; Lombard, 2014; Lorent et al., 2020). For example, sphingomyelin is known to be mainly restricted to the outer plasma membrane leaflet (Lorent et al., 2020). On the other hand, phosphoinositides and phosphatidylserine (PS) are found almost exclusively in the inner plasma membrane leaflet (Lorent et al., 2020). To preserve this membrane lipid transbilayer asymmetry, cells invest significant amounts of energy through ATP-dependent flippases and floppases that actively maintain the lipid distribution between the two leaflets of the plasma membrane (van Meer, 2011; Kobayashi and Menon, 2018). While the structure function relationship of nanodomains containing the molecular machineries for a diverse set of physiological functions has been studied in great detail, much less is known about the functional implications of the asymmetric transbilayer distribution of membrane lipids in biological membranes (Lorent et al., 2020; Doktorova et al., 2025). Nevertheless, it is widely assumed that this asymmetry is of broad relevance for proper cellular functions and that the disturbance of plasma membrane lipid transbilayer asymmetry can lead to malfunctions and potentially disease.

As of this day, only for a few cases, a causally determined relationship between membrane lipid transbilayer asymmetry and cellular functions has been established. An example is the regulation of the activity of membrane proteins depending on an asymmetric distribution of membrane lipids, being a critical parameter for cellular health (Pabst and Keller, 2024). Furthermore, membrane asymmetry of PS localized in the inner plasma membrane leaflet in benign cells is a prerequisite for signaling during apoptosis, a process that causes a global loss of membrane asymmetry by which PS is exposed on the cell surface and serves as an ‘eat me’ signal to phagocytes, removing deranged cells that may harm the entire organism (Kim et al., 2022). Another striking example is the temporary loss of plasma membrane lipid asymmetry that plays a critical role in maintaining normal physiological functions. For instance, recent studies revealed plasma membrane SM asymmetry to be disrupted during lysosomal damage. Calcium release from damaged lysosomes was shown to activate a SM scramblase, flipping SM to the cytosolic leaflet where it is converted into ceramide (Niekamp et al., 2022). This constitutes a signal that leads to lysosomal repair mediated by an ESCRT-independent pathway. Similarly, a loss of PE asymmetry has been observed during cytokinesis (Emoto et al., 1996). Although the mechanisms underlying this process are not yet fully understood, this phenomenon is crucial for effective cell division (Emoto et al., 2005). During cytokinesis, the two membrane leaflets are coupled, with PE being enriched in the outer leaflet of the cleavage furrow, alongside SM and cholesterol. By contrast, phosphatidylinositol PI(4,5)P_2_ is enriched in the inner leaflet. Disrupting the transbilayer distribution of any of these lipids was shown to impair cell division (Emoto and Umeda, 2000; Kunduri et al., 2022).

The strong coupling of membrane lipids such as SM, cholesterol, and PI(4,5)P_2_ between the two leaflets during cytokinesis shows intriguing similarities to the mechanism of FGF2 membrane translocation during unconventional secretion. Cholesterol has been demonstrated to tune PI(4,5)P_2_-dependent recruitment of FGF2 at the inner plasma membrane leaflet, resulting in substantial changes in the efficiency of unconventional secretion of FGF2 when plasma membrane cholesterol levels are manipulated (Lolicato and Nickel, 2022; Lolicato et al., 2022). Due to the intimate relationship between cholesterol and SM in the plasma membrane, it is likely that SM levels are also important for efficient secretion of FGF2. Along with the observation of Glypican-1 (GPC1), a GPI-anchored cell surface protein localized in cholesterol-rich ordered domains, being the primary heparan sulfate proteoglycan driving FGF2 secretion (Sparn et al., 2022a; Sparn et al., 2022b), the hypothesis of nanodomains containing the membrane translocation machinery for FGF2 was inspired (Lolicato and Nickel, 2022). In the current study, we now demonstrate that an asymmetric distribution of PI(4,5)P_2_ is a critical parameter for membrane pore formation mediated by PI(4,5)P_2_-dependent FGF2 oligomerization and to be essential for FGF2 secretion from cells. This observation therefore represents a striking example of the importance of transbilayer asymmetry of membrane lipids for a fundamental process in cell biology.

In terms of the molecular mechanism underlying FGF2 membrane translocation into the extracellular space, we propose the local accumulation of PI(4,5)P_2_ molecules underneath FGF2 oligomers recruited at the inner plasma membrane leaflet to produce a steep electrochemical gradient across the membrane that facilitates the opening of FGF2 induced lipidic membrane pores. Furthermore, with PI(4,5)P_2_ representing a non-bilayer lipid and due to its asymmetric distribution, we propose FGF2 oligomer-dependent accumulation of PI(4,5)P_2_ in a locally constrained manner to cause a massive amount of membrane stress. This in turn may compromise the structural integrity of the regular bilayer, facilitating the opening of a lipid membrane pore that is characterized by high membrane curvature, accommodating cone-shaped PI(4,5)P_2_ molecules. Thus, under these conditions, the opening of the pore reduces membrane stress. Following the capturing of FGF2 oligomers by GPC1 and their removal at the outer leaflet, the membrane pore may close spontaneously. This process has been imaged by single molecule TIRF microscopy in real time, demonstrating FGF2 membrane translocation events to be rather fast with average time intervals of about 200 ms (Dimou and Nickel, 2018; Dimou et al., 2019).

In conclusion, the findings presented in this study provide a compelling model, explaining membrane pore opening by FGF2 oligomer dependent clustering of PI(4,5)P_2_ molecules at the inner plasma membrane leaflet. Our results represent one of the few examples for a direct functional role of membrane lipid asymmetry for fundamental processes in cell biology and have general implications for the biophysical properties of biological membranes.

## Acknowledgements

We thank André Nadler (MPI-CBG Dresden, Germany) and Pavel Barahtjan, providing an expression construct encoding a fusion protein of a PH domain and the Halo tag as well as Hans-Michael Müller for a high-quality preparation of this fusion protein. This work was supported by the Deutsche Forschungsgemeinschaft (SFB/TRR 186, project A1; WN and FL, DFG Ni 423/10-1, DFG Ni 423/12-1 and DFG Ni 423/13-1; WN and DFG LO 2821/1-1; FL). The authors gratefully acknowledge the data storage service SDS@hd supported by the Ministry of Science, Research, and the Arts Baden-Württemberg (MWK), the German Research Foundation (DFG) through grant INST 35/1314-1 FUGG and INST 35/1503-1 FUGG.

## Materials and Methods

### Formation of Large Unilamellar Vesicles (LUVs)

**Table 1.** Lipid compositions used

| Lipid Composition (mol%) |
| --- |
| PC+2% or 3% or 4% or 5%PI(4,5)P<sub>2</sub>+1%RhPE |
| PC+5%PI(4)P+1%RhPE |
| PC+10%PE+5%PS+5%PI+30%CHO+15%SM+5%PI(4,5)P<sub>2</sub>+1%RhPE |
| PC+10%PE+5%PS+5%PI+30%CHO+15%SM+5%PI(4)P+1%RhPE |

\*Liss 16:0 RhPE is used as a fluorescent membrane marker

All the lipid were purchased from Avanti and stored in chloroform sealed with Argon at -20°C. Desired lipid concentrations were added to a 25mL round-bottom flask at final concentration of 4mM. Chloroform was evaporated under constant nitrogen flow while continuously shaking the flask to generate a thin lipid film. The lipid film was then hydrated using a buffer solution containing 10% (w/v) sucrose, 25mM HEPES-KOH, 150mM KCl pH 7.4, followed by vortexing to fully solubilize and hydrate the dried lipid film. To generate unilamellar vesicles, the solution was subjected to 10 freeze-thaw cycles, with each freeze (in liquid N2) and thaw (55°C water bath) step lasting 5 minutes. In the final step, the solution was subjected to 27 cycles of extrusion with 400nm filters. Liposomal size was measured by dynamic light scattering (observed size=200-400nm). The resulting vesicles were aliquoted and stored at -80°C for long-term storage. Each liposome aliquot was never freeze-thawed more than twice after its initial preparation.

### Formation of Giant Unilamellar Vesicles (GUVs)

**Table 2.** Lipid compositions used

| Lipid Composition (mol%) |
| --- |
| PC+5%PI(4,5)P<sub>2</sub>+1%Biotynl-PE+0.05%RhPE |
| PC+5%PI(4)P+1%Biotynl-PE+0.05%RhPE |
| PC+10%PE+5%PS+5%PI+30%CHO+15%SM+5%PI(4,5)P<sub>2</sub>+1%Biotynl-PE+0.05%RhPE |
| PC+10%PE+5%PS+5%PI+30%CHO+15%SM+5%PI(4)P+1%Biotynl-PE+0.05%RhPE |

\* Liss 16:0 RhPE is used as a fluorescent membrane marker
\* Biotynl-PE is used for immobilizing GUVs on Biotin-NeutrAvidin coated Ibidi chambers

The formation of GUVs was adapted from (Steringer et al., 2017) with minor modifications. Lipids at the desired concentration were mixed to prepare a working lipid mix solution. A total of 5 µL of this lipid mix was deposited on platinum electrodes (2.5 µL per electrode; Goodfellow Cambridge Limited 0.5 mm diameter, 99.99% pure, degree of hardening: glowed) Teflon Chambers were custom made by MPI Dresden/University Heidelberg). The electrodes were then dried for 10–15 minutes before submerging them in a 300 mM sucrose solution (301 Osmol). GUV formation was carried out for 50 minutes at 1.5V, 10Hz, 55°C. Following formation, GUVs were detached from the electrode surface for 25 minutes at 1.5V, 2Hz, 55°C. To prevent sudden temperature changes, before washing, the GUVs were gradually cooled to room temperature over 20 minutes. GUVs were then washed three times with 1200µL of 25mM HEPES-KOH, 150mM KCl pH 7.4 buffer, 306 Osmol (HK buffer) by centrifugation at 1200 rcf for 5 minutes at 25°C. After each wash, 1 mL of the supernatant was gently removed from the top, ensuring the lucid pellet remained undisturbed. In the final step, the pellet was resuspended and gently transferred to an 8 well Ibidi chambers.

The Ibidi chambers were pre-blocked with 0.1mg/mL Biotin-BSA for 15 minutes, followed by three washes with water, and was later blocked with 0.1mg/mL NeutrAvidin for 15 minutes, followed by three washes with water and one with HK buffer. Before adding 200 µL of washed GUVs to Ibidi chambers, 100 µL of HK buffer with Alexa 647 dye and His-FGF2-GFP or His-FGF2-Halo at a final concentration of 200nM was added. The protein was incubated with GUVs for approximately 1-1.5 hours before imaging for bulk quantification or for 15–20 minutes for kinetic measurement.

Confocal microscopy: Imaging was done with Zeiss LSM-800 using 1.4 oil immersion lens with 63x magnification. Two track-three channel was used for imaging, Channel 1-RhPE (561) (red-membrane marker to identify GUV) Channel 2-FGF2-GFP or FGF2-Halo-Alexa488 (488nm) (green-FGF2 fusion protein to analyse FGF2 membrane binding) Channel 3-Alexa647 (647nm) (gray-a small ≈1KDa dye which can enter lumen of the GUV in the event of pore formation). For all the conditions, GUVs with and without FGF2 fusion proteins were analysed. For recording time-series for FGF2-GFP translocation kinetics, after adding protein to the GUVs, time was marked as 0 min, GUV of interest was identified and imaged till 1.5-2 hours. Time-series was recorded with 30 secs, 5 mins or 10 mins difference in between the individual frames. After image acquisition, the data were processed and analyzed using Fiji either manually or by a macro script. Images were converted from .czi files into .tif and .jpg formats, with the latter used for visualization purposes.

For quantifying % of GUVs with pore, small circle was drawn inside the GUV and to the immediate outside of the GUV. If ratio of Alexa-647 fluorescence inside to the outside was equal to or above 0.6 mean threshold, GUV was declared to have the pore. For each experimental replica condition 20-59 GUVs were analyzed.

### Protein expression

Recombinant FGF2 fusion proteins and PI(4,5)P_2_ detector-PHx2-Halo were purified with E.coli strain BL21 Star using expression vector for His-FGF2-GFP, His-FGF2-Halo and His-PHx2-Halo. Protein expression was done with 2xYT media for overnight. All the proteins were purified on Ni-NTA affinity chromatography. For His-FGF2-GFP and His-FGF2-Halo, heparin chromatography was performed. In some case, additional, Superdex 75 column was also performed or PD-10 desalting column was done for buffer exchange. For PHx2-Halo protein, after initial Ni-NTA affinity chromatography, protein was subjected to HRV-3C protease to remove His tag. This was followed by Ni-NTA affinity chromatography to remove His-Tag. In the final step, Superdex 75 column was also performed to yield the pure protein. All the proteins were buffered with 25mM HEPES-KOH, 150mM KCl, pH=7.4 with 2% glycerol.

### FACS flow cytometry

FACS procedures were adapted from (Temmerman and Nickel, 2009). Prior to the experiment, Eppendorf tubes were blocked with fatty acid-free BSA unless stated otherwise. On the following day, BSA was removed and Eppendorf cups were washed with 25mM HEPES-KOH,150mM KCl pH 7.4 buffer (HK buffer). LUVs at a concentration of 1mM were washed in 500 µL HK buffer followed by incubation with 50µL of 2µM His-GFP, 2µM His-FGF2-GFP or 5µM His-Halo, 5µM His-FGF2-Halo for 1 hour, unless stated otherwise. Halo tag was detected using Halo ligand Alexa488. Unbound protein was washed with 500µL HK buffer by centrifugation for 10 minutes at 16000 rcf, 25°C. Supernatant containing unbound protein was removed carefully without disturbing the pellet. In the final step, pellet was resuspended in 500µL HK buffer unless stated otherwise. For flow cytometry, liposomes were gated based on size—light scattering—and rhodamine-derived fluorescence—incorporated in the liposomal membrane. Minor background signals from liposomes incubated with GFP or Halo +Alexa488 were subtracted from the signals of liposomes incubated with His-FGF2-GFP or His-FGF2-Halo +Alexa488. The resulting signal was then normalized to 5 mol% PI(4,5)P_2_ vesicles.

### Sedimentation assay

1mM LUVs were washed in 100µL HK buffer, followed by incubation with 50µL of 4µM His-FGF2-GFP or 5µM His-FGF2-Halo for 1 hour or 3 hours. The vesicles were then centrifuged at 16000 rcf for 10 minutes at 25°C, and the unbound FGF2 was collected as the supernatant. The pellet, containing vesicles bound to the FGF2 fusion protein, was washed twice with 100µL HK buffer, and the supernatant was discarded. In the final step, the pellet (containing vesicles bound to the protein) were resuspended in 1× reducing SDS-sample buffer. Both the unbound and bound fractions of the FGF2 fusion protein were analyzed on a 1.5mm, 4–12% Bis-Tris SDS-PAGE in MES buffer. Proteins were stained with Coomassie instant blue. For analysis, the gel was scanned using an EPSON Perfection 4870 photo scanner or a LICOR Odyssey system. For analysis, FGF2 bound to the 5%PI(4,5)P_2_ vesicles, was set to 100% and data was normalised accordingly.

### Kinase reaction to generate asymmetric LUVs/GUVs

Both LUVs and GUVs were subjected to kinase reaction to generate PI(4,5)P_2_ asymmetric vesicles. In the first step, GUVs or 1mM LUVs were washed with 25mM HEPES-KOH, 150mM KCl pH 7.4 (HK buffer). To the liposome pellet, premixed reaction ingredients 50mM TRIS-HCl (pH=7,4), 5mM MgCl_2_, 100µM ATP, 20nM PIP5K1C in HK buffer were added (1mM liposome≈50µM PI(4)P or PI(4,5)P_2_). Osmolarity of luminal and extra-luminal solution do not differ by more than 10-20 Osmol. Reaction was carried out at 25°C for 15mins with shaking at 600rpm for LUVs and 350rpm for GUVs. Liposomes were washed twice, with 50-500µL HK buffer for LUVs (depending on the assay) and 1mL HK buffer for GUVs to remove all the unreacted ingredients or damaged vesicles. Following the reaction, vesicles were analyzed with sedimentation assay, flow cytometry, and confocal imaging.

### Testing lipid asymmetry in LUVs

PC+5%PI(4,5)P_2_ (positive control for FGF2 binding) and PC+5%PI(4)P (substrate liposome) vesicles were used as controls. For kinase reaction 50mM TRIS HCl, 5mM MgCl_2_, 100µM ATP, 50µM PI(4)P (1mM total concentration in liposome), and 20nM PIP5K1C was used. As additional controls, reaction ingredients were added to the PI(4,5)P_2_ vesicles and mock was performed with PI(4)P vesicles. After completion of the reaction, reaction mixture was suspended in 50µL of 25mM HEPES-KOH, 150mM KCl pH 7.4 (HK buffer) and centrifuged at 16000 rcf, 10 mins at 25°C. Supernatant was discarded and the washing step was repeated once more.

For non-treated samples, post washing, the pellet was resuspended in 50µL of 1µM FGF2-GFP or 1µM GFP for 30 mins. After incubation, 50µL of HK buffer was added, and the sample was centrifuged at 16000 rcf for 10 mins at 25°C. The supernatant was discarded, and the washing step was repeated one more time. In the final step, the pellet was resuspended in 350µL of HK buffer for FACS measurement. (Note: the pellet can be stubborn when binding occurs, so it is important to resuspend it a minimum of 50-100 times.)

For reformation, the washed pelleted vesicles were disintegrated with 50µL HK buffer and 100µL of a 40% MeOH/CHCl₃ solution using 5 seconds of vortexing. The organic and aqueous layers were allowed to separate for 2-5 mins. The organic layer was carefully collected using a pipette (the tip should be pre-filled with the 40% MeOH/CHCl₃ solution to avoid sucking in the aqueous phase). The aqueous layer was washed twice more with 100 µL of 40% MeOH/CHCl₃ to ensure complete transfer of lipid content to the organic layer. The organic layer was then dried by purging with nitrogen gas and was placed under vacuum for at least 2 hours. The dried lipid film was hydrated with 50µL of 10% (w/v) sucrose HK buffer. To generate unilamellar vesicles, sample was subjected to 10 freeze-thaw cycles. After this stage, 5µL is sample was collected for dynamic light scattering measurement. The sample was then diluted with 100µL of HK buffer and centrifuged at 16000 rcf for 10 mins at 25°C. After centrifugation, the pellet was resuspended in 50µL of 1 µM FGF2-GFP or 1 µM GFP for 30 mins. After incubation, 50 µL of HK buffer was added, and the sample was centrifuged at 16000 rcf for 10 mins at 25°C. The supernatant was discarded, and the washing step was repeated one more time. In the final step, the pellet was resuspended in 350 µL of HK buffer for FACS measurement. FACS flow cytometry measurements were performed for all liposome samples containing GFP or FGF2-GFP. For analysis, FGF2-GFP bound to the 5%PI(4,5)P_2_ vesicles, was set to 100% and data was normalised accordingly.

### Testing lipid asymmetry in GUVs

GUVs with PM+5mol% PI(4,5)P_2_ and PM+5mol% PI(4)P were prepared, washed (once) and immobilized on glass bottom ibidi chambers as described previously in this section. Vesicle was identified using Zeiss LSM-800 using 1.4 oil immersion lens with 63x magnification. Imaging involved two channels: Channel 1-RhPE (561) (red-membrane marker to identify GUV) Channel 2-Alexa647 (647nm) (gray-a small ≈1KDa dye which can enter lumen of the GUV in the event of pore formation). To the GUVs Mg^2+^, ATP, PIP5K1C (for concentrations, see “*Kinase reaction to generate asymmetric LUVs/GUVs*” section) along with tracer dye Alexa647 was added and time was marked as 0mins. Vesicles was monitored for 15minutes and time series was recorded with 1 min difference in each individual frame.

### Culturing cells CHO-K1

CHO-K1 cells, expressing FGF2-GFP in a doxycycline dependent manner were used. The cells were cultured at 37°C in αMEM media. They were maintained for 20 passages or approximately two months after thawing. Before the experiment FGF2-GFP expression was induced with doxycycline for overnight.

### Disrupting transbilayer PI(4,5)P_2_ and PS asymmetry in cells

Approximately, 10000 cells were seeded in 8 well ibidi chambers with or without 1:1000 doxycycline dilution, overnight prior to experiment. Lipids at the desired concentration were placed in an Eppendorf tube and dried for approximately 1 hour under vacuum. They were then resuspended in a 6mM EDTA solution and processed with the following cycle: 30 seconds of vortexing, 5 minutes in a 55°C water bath, 30 seconds of vortexing, and 5 minutes of sonication. This cycle was repeated once more. Micelle/vesicle size was measured using dynamic light scattering (observed size=multi-populational distribution below and above 100nm).

PI(4,5)P_2_ treatment: Cells were treated with 15nM lipid in 1.5mM EDTA using FCS-free media, with a final volume of 50µL lipid EDTA solution in 150µL FCS-free media per chamber. The cells were incubated at 37°C for 20 minutes.

Mock treatment: Cells were treated with 1.5mM EDTA using FCS-free media, with a final volume of 50µL of EDTA solution in 150µL FCS-free media per chamber. The cells were incubated at 37°C for 20 minutes.

No treatment: Cells were not treated with any solution.

*[Calculation description: For PI(4,5)P_2_ - Total lipids in PM assumed to be 10^9^ lipids, assuming 3% PI(4,5)P_2_ in the PM, three times more PI(4,5)P_2_ was added in the experiment. EDTA was added because of Ca^2+^ and Mg^2+^ in the media. Divalent cations aggregate PI(4,5)P_2_, which hinders the incorporation of PI(4,5)P_2_ in the outer leaflet. Total Ca^2+^+ Mg^2+^ in media is approximately ≈1,5mM, so 1.5mM EDTA was used in the experiment. For PS lipid - It is used as a control lipid, and was added similar to PI(4,5)P_2_]*

After each treatment, cells were washed with PBS three times. After this, either cells were fixed with 3%PFA for 13mins at r.t. or were kept live in LCI solution.

For detecting PI(4,5)P_2_, 40nm PHx2-Halo –detected with Alexa fluor 660—was used for 25-30mins at r.t. Before imaging, cells were washed three times with PBS to remove unbound protein and were kept in PBS (for fixed cell imaging) or LCI (for live cell imaging). For PS detection, only live cells could be used, as fixing of cells led to PS exposure on the exoplasmic leaflet. AnnexinV-Alexa647 with 2mM Ca^2+^ in LCI solution was used as PS detector protein at 1.5:100 dilution. Imaging was done instantly after addition of AnnexinV to cells.

For time course study, the goal was to observe retention of PS and PI(4,5)P_2_ on the outer leaflet of PM over time. After disturbing the PI(4,5)P_2_ lipid, cells were fixed at either 0mins, 20mins, 30mins, 40mins, 60mins and analyzed with PHx2-Halo-Alexa fluor 660.

Imaging and processing: Cells were imaged using Zeiss LSM-800 using 1.4 oil immersion lens with 63x magnification. Two track channel was used for imaging, Channel 1-PHx2-Halo detection with Alexa660 (660) or AnnexinV-Alexa647 (647nm) (white) and Channel 2-FGF2-GFP (488nm) (green). After image acquisition, the data were processed and analyzed using Fiji. A macro script was used to convert the .czi files from the LSM800 microscope into .tif and .jpg formats, with the latter used for visualization purposes.

### FGF2 Translocation assay using confocal microscope

Approximately 10000 cells were seeded with 1.1:1000 doxycycline, overnight prior to the experiment. On the following day, cells were washed with PBS three time. Cells were either left untreated, mock-treated, or treated with PI(4,5)P_2_ as described above. Following the treatment, cells were washed three times with PBS, then incubated with 1 mg/mL heparin for 10 minutes at room temperature. For effective heparin wash, indicated cell density was crucial. After another three PBS washes, cells were placed in LCI for 0, 10, 20, 30, 40, or 60 minutes, followed by fixation with 3% PFA for 13 minutes at room temperature. To visualize surface FGF2-GFP, cells were incubated with an anti-GFP-Alexa647 antibody at 1:400 dilution in PBS for 30 minutes at room temperature. This was followed by three washes with PBS, after which the cells were stored in PBS and imaged using confocal microscopy.

Imaging and quantification: Cells were imaged using Zeiss LSM-800 using 1.4 oil immersion lens with 63x magnification. Two track channel was used for imaging, Channel 1-AntiGFP-Alexa647 (647nm) (white) and Channel 2-FGF2-GFP (488nm) (green). For each condition, 15–40 cells were imaged.

After image acquisition, the data were processed and analyzed using Fiji. A macro script was used to convert the .czi files from the LSM800 microscope into .tif and .jpg formats, with the latter used for visualization purposes. For quantifying surface FGF2-GFP—detected via antiGFP-Alexa647—the macro script was applied to the .tif files. After setting an appropriate threshold, individual cells in each image were identified and saved as Regions of Interest (ROIs). A band was drawn near the cell boundary to account for the cellular membrane, and the intensity of anti-GFP-Alexa647 was recorded. Each ROI was manually inspected to ensure accurate cell selection. In cases of misidentification, the ROI was adjusted and updated accordingly. As a final step, the “surface FGF2-GFP after heparin wash at 0 minutes” condition was subtracted from all-time points to account for nonspecific background. The processed data were plotted in Prism, and statistical analysis was performed using a one-way ANOVA (Brown-Forsythe and Welch ANOVA test) with Dunnett T3 (n>50 per group) test.

### Materials

**Table 3.** Lipid compositions used for vesicles preparation

| Lipid | Order number | Company |
| --- | --- | --- |
| Liver PC | 840055P | Sigma Avanti |
| Liver PE | 840026P/840026C | Sigma Avanti |
| Brain PS | 840032P/840032C | Sigma Avanti |
| Liver PI | 840042P/840042C | Sigma Avanti |
| Brain PI(4,5)P2 | 840046P | Sigma Avanti |
| Brain PI(4)P | 840045P | Sigma Avanti |
| Egg SM | 860061P | Sigma Avanti |
| Cholesterol | 700000P | Sigma Avanti |
| BiotynIPE | 870282P/870282C | Sigma Avanti |
| 16:0Lis RhPE | 810158P | Sigma Avanti |

\*All the lipid were purchased from avanti and stored under Argon

**Table 4.** Reagents used in this study

| Reagent/Protein | Order number | Company |
| --- | --- | --- |
| LC-Heparin | H3149-10KU | Sigma |
| Sucrose | 4621.1 | Roth |
| Biotin-BSA | A8549 | Sigma |
| Neutravidin | A2666 | Invitrogen/ThermoFisher |
| Alexa647 | A33084 | Invitrogen/ThermoFisher |
| PIP5K1C | A33522 | ThermoFisher |
| MgCl<sub>2</sub> | 63068 | Fluka |
| CaCl<sub>2</sub> | 31307 | Sigma |
| HEPES | H4034 | Sigma |
| KCl | SZBG3080H | Fluka |
| KOH | 30603 | Sigma |
| ATP | 10 519 979 001 | Roche |
| EDTA | 1.08418.1000 | Sigma |
| BSA fatty free | REF 10775 835 001 | Roche |
| Paraformaldehyde 16%<br>solution (PFA) | 15210 | Electron Microscopy<br>Science |
| Alpha-MEM | M8042 | Sigma |
| Dulbecco PBS | D8537 | Sigma |
| Live cells imaging solution<br>(LCI) | A14291DJ | ThermoFisher |
| Halo Ligand-Alexa Fluor<br>660 | G847B | Promega |
| Halo Ligand-Alexa Fluor<br>488 | G1001 | Promega |
| AntiGFP-Alexa647 | A-31852 | ThermoFisher |
| PHX2-Halo-Halo | - |  |
| His-FGF2-GFP | - |  |
| His-FGF2-Halo | - |  |
| AnnexinV-Alexa647 | A23204 | ThermoFisher |

***Video 1***

PM+5mol%PI(4,5)P_2_ GUV treated with Mg^2+^, ATP, PIP5K1C and tracer dye Alexa-647 for 15mins: Giant unilamellar vesicles (GUVs) were prepared with 5 mol% PI(4,5)P₂ along with membrane marker Rhodamine-PE, in PM like background. After immobilizing the GUVs, Mg^2+^, ATP, PIP5K1C and small trace dye Alexa-647 was added and time was marked as 0 min. Vesicle was monitored for the entire course of kinase reaction i.e. 15mins.

***Video 2***

PM+5mol%PI(4)P GUV treated with Mg^2+^, ATP, PIP5K1C and tracer dye Alexa-647 for 15mins: Giant unilamellar vesicles (GUVs) were prepared with 5 mol% PI(4)P along with membrane marker Rhodamine-PE, in PM like background. After immobilizing the GUVs, Mg^2+^, ATP, PIP5K1C and small trace dye Alexa-647 was added and time was marked as 0 min. Vesicle was monitored for the entire course of kinase reaction i.e. 15mins.

***Video 3***

FGF2 translocation kinetics for symmetric PM+5mol% PI(4,5)P_2_ GUV in real time: Giant unilamellar vesicles (GUVs) were prepared with 5 mol% PI(4,5)P₂ along with membrane marker Rhodamine-PE, in PM like background. Small trace dye Alexa-647 was added to record the event of pore formation for kinetic measurement. After addition of FGF2-GFP, time was marked as 0 min and vesicle was given 10-20 mins to immobilize before starting the time series.

***Video 4***

FGF2 translocation kinetics for asymmetric PM+5mol% PI(4,5)P_2_ GUV in real time: Giant unilamellar vesicles (GUVs) were prepared with 5 mol% PI(4)P along with membrane marker Rhodamine-PE, in PM like background. Vesicles were subjected to kinase reaction with Mg^2+^ + ATP + PIP5K1C for conversion of PI(4)P to PI(4,5)P₂ to yield asymmetric vesicle. Small trace dye Alexa-647 was added to record the event of pore formation for kinetic measurement. After addition of FGF2-GFP, time was marked as 0 mins and vesicle was given 10-20 mins to immobilize before starting the time series.

***Video 5:***

FGF2 translocation kinetics for symmetric PM+5mol% PI(4,5)P_2_ GUV in real time: Giant unilamellar vesicles (GUVs) were prepared with 5 mol% PI(4,5)P₂ along with membrane marker Rhodamine-PE, in PM like background. Vesicles were treated with Mg^2+^ + ATP + PIP5K1C as a control. Small trace dye Alexa-647 was added to record the event of pore formation for kinetic measurement. After addition of FGF2-GFP, time was marked as 0 mins and vesicle was given 10-20 mins to immobilize before starting the time series. In the presented video, the time series starts at 23 mins. However, since the vesicle was not immobilized until 33 mins, the cropped video excludes the 23–33 minute segment. Please refer to the uncropped video for the full sequence.

